# SARS-CoV-2 Variant-Specific Differences in Inhibiting the Effects of the PKR-Activated Integrated Stress Response

**DOI:** 10.1101/2022.12.21.521388

**Authors:** Wanda Christ, Jonas Klingström, Janne Tynell

## Abstract

The integrated stress response (ISR) is a eukaryotic cell pathway that triggers translational arrest and the formation of stress granules (SGs) in response to various stress signals, including those caused by viral infections. The SARS-CoV-2 nucleocapsid protein has been shown to disrupt SGs, but SARS-CoV-2 interactions with other components of the pathway remains poorly characterized. Here, we show that SARS-CoV-2 infection triggers the ISR through activation of the eIF2α-kinase PKR while inhibiting a variety of downstream effects. In line with previous studies, SG formation was efficiently inhibited and the induced eIF2α phosphorylation only minimally contributed to the translational arrest observed in infected cells. Despite ISR activation and translational arrest, expression of the stress-responsive transcripts ATF4 and CHOP was not induced in SARS-CoV-2 infected cells. Finally, we found variant-specific differences in the activation of the ISR between ancestral SARS-CoV-2 and the Delta and Omicron BA.1 variants in that Delta infection induced weaker PKR activation while Omicron infection induced higher levels of p-eIF2α and greatly increased SG formation compared to the other variants. Our results suggest that different SARS-CoV-2 variants can affect normal cell functions differently, which can have an impact on pathogenesis and treatment strategies.

## Introduction

Coronaviruses are a family of enveloped, positive-sensed, single-stranded RNA viruses within the *Nidovirales* order. They infect a variety of animals like birds, bats and other mammals including humans [1]. Since its emergence in November 2019, the *betacoronavirus* severe acute respiratory syndrome coronavirus 2 (SARS-CoV-2) has quickly spread around the world causing coronavirus disease 2019 (COVID-19) [2–4]. SARS-CoV-2 has mutated over time, giving rise to several variants associated with higher infectivity and increased escape from neutralizing antibodies [5-6]. This includes Delta (B.1.617.2) which was first described in India in October 2020 and Omicron (B.1.1.529) which was first described in South Africa in November 2021 [7-8]. The latter is still dominating in the population to this date [9].

Viral infections induce a variety of stress stimuli within the host cells which can trigger stress sensors and thereby activate the integrated stress response (ISR). Activation of the ISR-pathway allows the cells to respond to the stress, either by resolving it or by inducing apoptosis [10-11]. The ISR is activated by phosphorylation of the alpha subunit of eukaryotic initiation factor 2 (eIF2α) by four distinct kinases each sensing a different type of stress. Heme regulated eIF2α-kinase (HRI, EIF2AK1) gets activated during oxidative stress. dsRNA-activated protein kinase R (PKR, EIF2AK2) senses double-stranded RNA (dsRNA), PKR-like kinase (PERK, EIF2AK3) detects misfolded proteins in the endoplasmic reticulum and general control nonderepressible 2 (GCN2, EIF2AK4) senses amino acid availability and responds to a lack of amino acids as well as glucose deprivation [11-13]. Phosphorylation of eIF2α, by the above mentioned kinases, triggers a signaling pathway leading to a reduction in global translational levels and an increased expression of a few selected genes (stress-responsive transcripts) that enable the cell to recover from the stress and terminate the ISR [14]. As a result of the translational shutdown, stalled initiation complexes containing silenced mRNAs, small ribosomal subunits, translation initiation factors and other RNA-binding proteins accumulate within the cytoplasm to form stress granules (SGs) [15-16]. These structures are thought to function as a storage for the initiation complexes until translation initiation is restored. SGs also provide certain antiviral functions, for example by recruiting and activating antiviral proteins or by sequestering viral factors [10, 17]. Viruses are dependent on the host cell translation machinery for their own protein production and therefore many viruses have developed strategies to interfere with the ISR and SG formation [18]. A common approach, also employed by other coronaviruses like MERS-CoV and infectious bronchitis virus (IBV), is to inhibit the eIF2α-kinase PKR [19-20]. Other viruses inhibit eIF2α-phosphorylation or cleave G3BP, a protein that is essential for SG formation [21-22]. The SARS-CoV-2 nucleocapsid protein (N protein) has been shown to inhibit SG formation via binding to G3BP1 and G3BP2 [23–25]. However, many questions about the activation and role of the ISR during SARS-CoV-2 infection remain open. Here, we show that the ISR is activated during SARS-CoV-2 infection via activation of PKR, but despite phosphorylation of eIF2α only limited SG formation or translation of stress-responsive transcripts is observed, indicating SARS-CoV-2 mediated active inhibition of the ISR pathway. Chemical stimulation of the stress kinases shows viral resistance especially towards stress mediated through PKR but also a general inhibition of stress-responsive transcript translation. Finally, we show specific differences in the activation and inhibition of the ISR pathway between the ancestral strain and the Delta and Omicron variants, highlighting SARS-CoV-2 sub lineage differences in manipulation of cell responses.

## Results

### SARS-CoV-2 infection activates the integrated stress response through PKR but does not trigger widespread SG formation

To investigate whether the ISR gets activated during SARS-CoV-2 infection, levels of phosphorylated eIF2α (p-eIF2α) were measured in Vero E6 cells, an African green monkey kidney cell line, and in A549-hACE2 cells, a human lung epithelial cell line overexpressing human ACE2. At 24 hours post-infection (hpi), phosphorylation of eIF2α was detected in both cell lines (Fig. 1A). The eIF2α-kinases PKR, PERK and GCN2 are known to be activated upon infections with various viruses [26]. To analyze their activation during SARS-CoV-2 infection, we examined their phosphorylation status and expression levels in A549-hACE2 cells. As shown in an earlier study [27], strong phosphorylation of PKR was observed, while levels of p-PERK remained unchanged in infected cells (Fig. 1B and 1C). For GCN2, phosphorylation was not detected but a significant reduction of the protein was observed in infected cells, an effect that has also been previously shown during infection with SARS-CoV-1 [28] (Fig. 1B and 1C).

**Figure 1:**
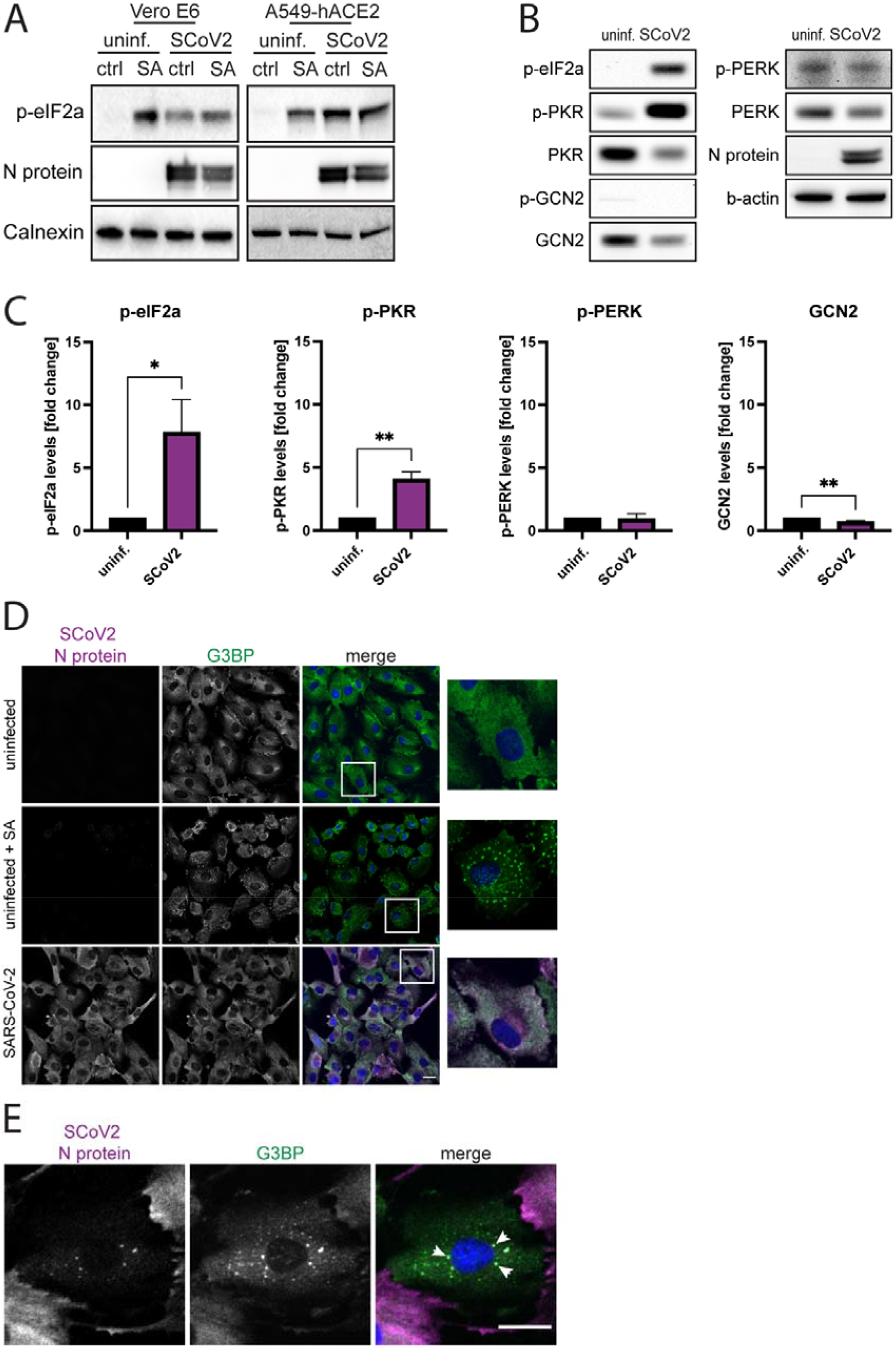
ISR activation and SG formation in ancestral SARS-CoV-2 infected cells. (A) eIF2α-phosphorylation in SARS-CoV-2 infected Vero E6 and A549-hACE2 cells. Uninfected and SARS-CoV-2 infected cells were treated with 1mM sodium arsenite (SA), an ISR-inducing chemical, for 1h, as positive control. Cell lysates were collected at 24 hpi and analyzed for levels of p-eIF2α. Calnexin was used as a loading control. (B) Protein levels and phosphorylation of eIF2α-kinases in SARS-CoV-2 infected A549-hACE2 cells. Cells were infected SARS-CoV-2. Cell lysates were collected at 24 hpi and analyzed for expression and phosphorylation levels of different eIF2α-kinases. β-actin was used as a loading control. (C) Quantification of (B). Protein levels were normalized to cellular levels of β-actin. The data are presented as the fold change in relation to the protein levels of uninfected cells. Data are represented as mean ± SEM. n = 3. (D) G3BP staining in SARS-CoV-2 infected A549-hACE2 cells. Cells were infected with SARS-CoV-2 and fixed after 24h. Uninfected cells that were either untreated or treated with 1mM SA were used as a control. The cells were stained with antibodies against SARS-CoV-2 N protein (magenta) and G3BP (green) and with DAPI (blue). Imaging was performed with confocal microscopy (20X). Scale bars 20 μm. (E) Localization of the viral N protein within SGs. A549-hACE2 cells were fixed 24h hpi and stained with antibodies against SARS-CoV-2 N protein (magenta) and G3BP (green) and with DAPI (blue). Arrows show colocalization of viral proteins with G3BP. Imaging was performed with confocal microscopy (20X). Scale bars 20 μm. *, p < 0.05; **, p < 0.005

To assess whether the eIF2α-phosphorylation was associated to formation of SGs, infected cells were stained for the SG marker G3BP1 and inspected via immunofluorescence. As reported by others [24], the vast majority of SARS-CoV-2 infected cells were negative for SGs (Fig. 1D, Supp. Fig. 1). The few infected cells positive for SGs showed lower levels of N protein and co-localization of N protein with G3BP within SGs (Fig. 1E, Supp. Fig. 1, Supp. Fig. 3).

Together, these results show that while the ISR is activated during SARS-CoV-2 infection, as indicated by phosphorylation of eIF2α and PKR, this does not trigger widespread SG formation.

### SARS-CoV-2 infected cells are resistant to oxidative and PKR-mediated stress

The SARS-CoV-2 N protein has been shown to inhibit SG formation by binding to the SG-nucleating protein G3BP [23-24]. As this occurs downstream of eIF2α-phosphorylation the N protein can therefore potentially inhibit all types of ISR-activating stress signals. To study the ability of SARS-CoV-2 to inhibit SG formation in more detail, uninfected and infected Vero E6 cells were exposed to the stress-inducers sodium arsenite, Poly(I:C), thapsigargin, or starvation, and then examined for SG formation (Fig. 2A). As previously reported, SG formation was inhibited in SARS-CoV-2 infected cells exposed to sodium arsenite, an inducer of oxidative stress [23-24] (Fig. 2B). A reduction in SG formation in infected compared to uninfected cells was also observed after Poly(I:C) treatment, which triggers activation of PKR (Fig. 2C). The ER stress inducer thapsigargin and starvation both induced SG-formation in infected cells but not in uninfected cells (Fig. 2D and 2E). Together, this suggests that SARS-CoV-2 can inhibit SG-formation triggered by oxidative stress and dsRNA, but infected cells remain susceptible and even sensitized towards SG formation induced via ER stress and starvation.

**Figure 2:**
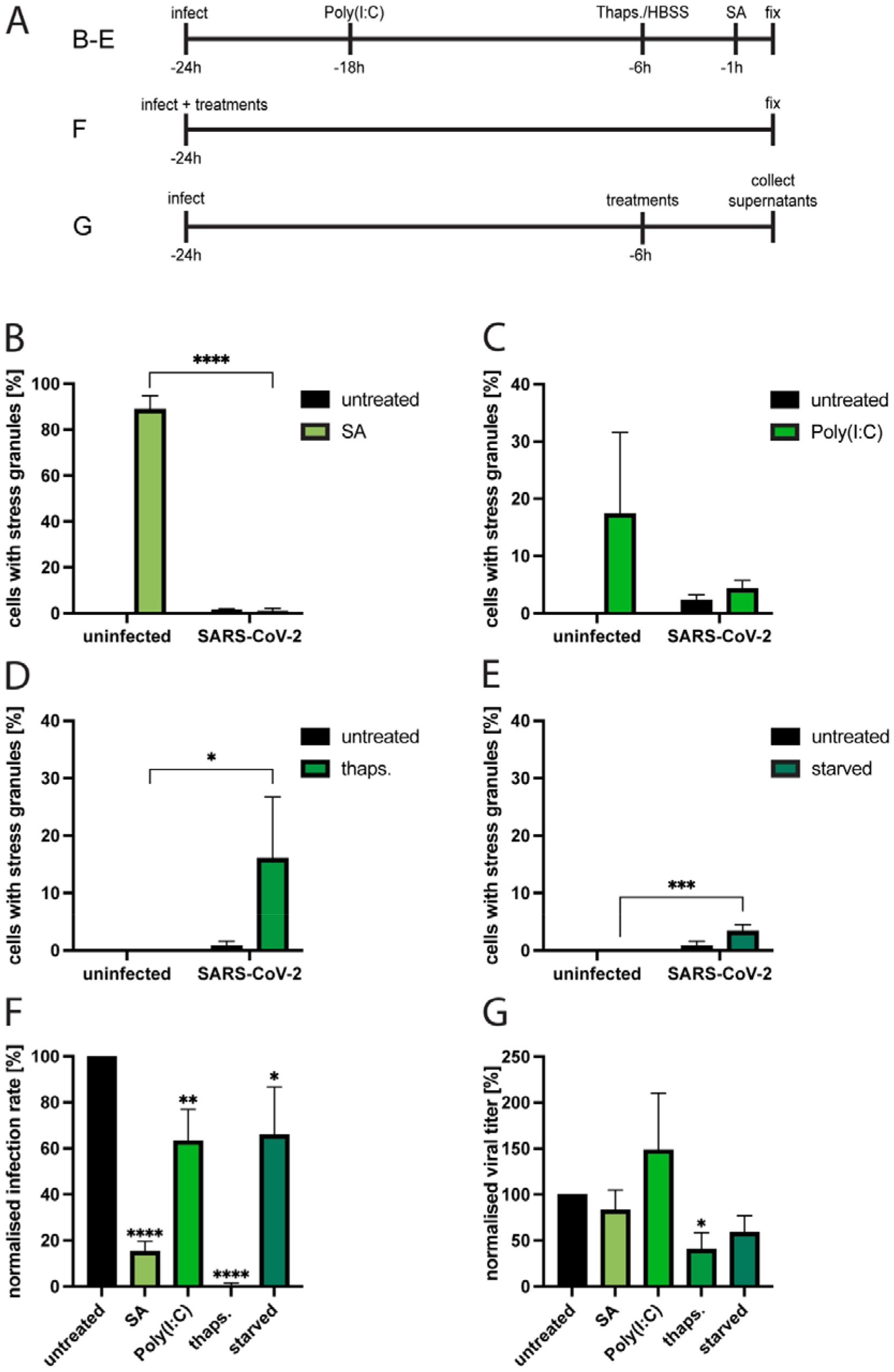
The effect of different stress stimuli on SG formation during ancestral SARS-CoV-2 infection and on viral replication in Vero E6 cells. (A) Experimental setup for figures 2B -G. Uninfected and ancestral SARS-CoV-2 infected cells were (B) treated with 1mM sodium arsenite for 1h, (C) transfected with 2.5μg Poly(I:C) overnight, (D) treated with 5μM thapsigargin for 5h or (E) starved for 5h and fixed at 24 hpi. They were stained for SARS-CoV-2 N protein and G3BP. For the analysis using fluorescence microscopy, ≥5 images of a total of >100 cells were taken at random positions and the number of infected cells showing SGs was determined. Data are represented as mean ± SEM. n ≥ 3. (B) Effect of different stress inducers on infectivity. SARS-CoV-2 infected cells were treated with sodium arsenite, Poly(I:C), thapsigargin or starved (see materials and methods) and fixed at 24 hpi They were stained for SARS-CoV-2 N protein. For the analysis using fluorescence microscopy, ≥5 images of a total of >100 cells were taken at random positions and the number of infected cells was determined and normalized against the infection rate of untreated cells. Data are represented as mean ± SEM. n = 3. (C) Effect of different stress inducers on viral replication. SARS-CoV-2 infected cells were treated with sodium arsenite, Poly(I:C), thapsigargin or starved (see materials and methods). 6h after treatment the viral titers were determined and normalized against the titers in untreated cells. Data are represented as mean ± SEM. n ≥ 3. *, p < 0.05; **, p < 0.005; ****, p < 0.0001

To better understand the effects of different stress stimuli on SARS-CoV2 infectivity, we investigated the infection rate in Vero E6 cells when adding the virus together with sodium arsenite, Poly(I:C) or thapsigargin or with starvation medium (Fig. 2A). All treatments negatively affected the infection rate, and the strongest effect was observed for thapsigargin, which almost completely inhibited infection when administered with the virus (Figure 2F). To analyze the effect of the treatments on viral replication, infected cells were first infected, and at 18 hours hpi treated with the different stress inducers for 6 hours before collection of supernatants at 24 hpi. Viral titers were then analyzed in the supernatants (Figure 2A). Again, thapsigargin had the largest impact on progeny virus production, but interestingly sodium arsenite barely affected it and Poly(I:C) might even promote progeny virus production (Fig. 2G). This indicates that while SARS-CoV-2 is susceptible towards the cellular stress response during the initial infection step, progeny virus production is not strongly decreased in infected cells where stress responses are induced.

### The SARS-CoV-2 ancestral strain, Delta, and Omicron BA.1 show differences in ISR activation and SG formation

The SARS-CoV-2 variants Delta and Omicron have accumulated large numbers of mutations throughout their genome [29]. While mutations of the spike protein are known to impact receptor binding efficiency and neutralizing antibody escape, little is known about possible differences in intracellular virus-host interactions between different SARS-CoV-2 variants. To investigate possible variant specific differences in suppressing SG formation, ancestral SARS-CoV-2, Delta, and Omicron BA.1 infected Vero E6 cells were analyzed for SG formation at different time points post-infection. The SARS-CoV-2 ancestral strain and Delta showed a similar pattern with a low prevalence of SGs throughout the infection (Fig. 3A). However, Omicron induced significantly higher levels of SG-positive cells than the other variants at 24 hpi and 48 hpi (Fig. 3A). This difference was even more pronounced in A549-hACE2 cells, where around 5% of ancestral strain or Delta infected cells were SG-positive compared to almost 60% of Omicron infected cells (Fig. 3B).

**Figure 3:**
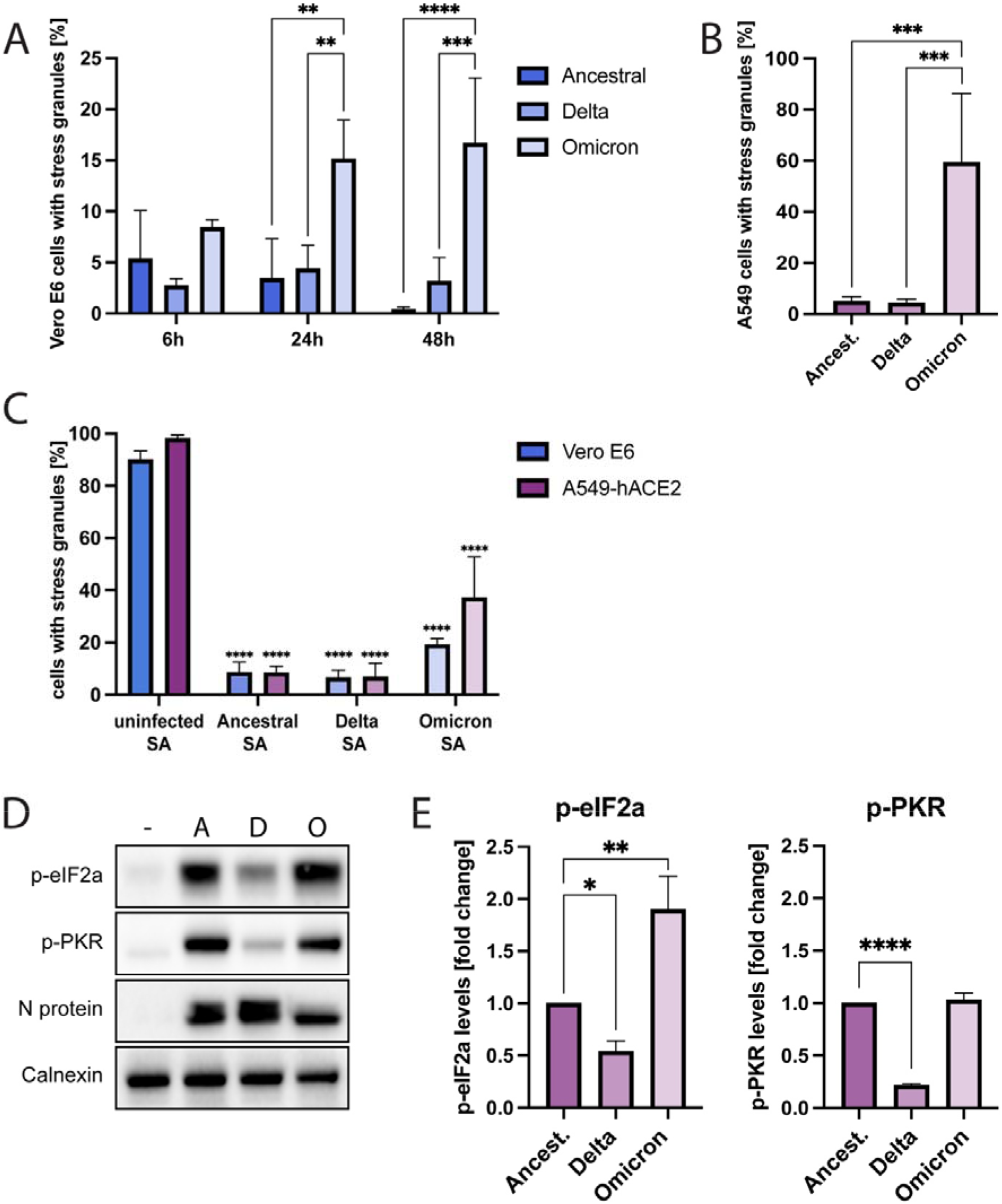
ISR activation and SG formation during infection with different SARS-CoV-2 variants. (A) Time kinetics of SG formation in SARS-CoV-2 infected Vero E6 cells. Cells infected with different SARS-CoV-2 variants were fixed at 6, 24 or 48 hpi. They were stained for SARS-CoV-2 N protein and G3BP. For the analysis using fluorescence microscopy, ≥5 images of a total of >100 cells were taken at random positions and the number of infected cells showing SGs was determined. Data are represented as mean ± SEM. n ≥ 3. (B) SG formation in SARS-CoV-2 infected A549-hACE2 cells. Cells infected with different SARS-CoV-2 variants were fixed at 24 hpi. They were stained for SARS-CoV-2 N protein and G3BP. For the analysis using fluorescence microscopy, ≥5 images of a total of >100 cells were taken at random positions and the number of infected cells showing SGs was determined. Data are represented as mean ± SEM. n ≥ 3. (C) SG formation after sodium arsenite challenge in SARS-CoV-2 infected Vero E6 and A549-hACE2 cells. Cells were infected with different SARS-CoV-2 variants and treated with 1mM sodium arsenite for 1h. Cells were fixed at 24 hpi. They were stained for SARS-CoV-2 N protein and G3BP. For the analysis using fluorescence microscopy, ≥5 images of a total of >100 cells were taken at random positions and the number of infected cells showing SGs was determined. Data are represented as mean ± SEM. n = 3. (D) eIF2α- and PKR-phosphorylation in SARS-CoV-2 infected A549-hACE2 cells. Cells were infected with ancestral (A), Delta (D), or Omicron (O) SARS-CoV-2. Lysates were collected at 24 hpi and analyzed for levels of p-eIF2α and p-PKR. Calnexin was used as a loading control. (E) Quantification of (D). Protein levels were normalized to cellular levels of β-actin. The data are presented as the fold change in relation to the protein levels of cells infected with ancestral SARS-CoV-2. Data are represented as mean ± SEM. n = 3. *, p < 0.05; **, p < 0.005; ***, p < 0.0005; ****, p < 0.0001

As we observed higher frequency of SG-positive cells after Omicron-infection, we next analyzed for possible variant-specific differences in capacity to suppress SG-formation. All three variants efficiently suppressed sodium arsenite-induced SG formation in both cell types (Fig. 3C), suggesting a common mechanism to inhibit oxidative stress mediated SG-formation among SARS-CoV-2 variants. The N protein has been suggested to be important for SARS-CoV-2 mediated inhibition of SGs [23]. In line with this, lower N protein levels were observed in ancestral strain and Delta infected SG-positive, compared to SG-negative, cells (Suppl. Fig. 2A and 2B). In contrast, no difference in N protein levels were observed between SG-positive and SG-negative omicron-infected cells (Suppl. Fig. 2A and 2B). This suggests that the Omicron N protein might be less effective in inhibiting SG formation which could explain why SGs appear more frequently in Omicron-infected cells.

To further investigate if the observed differences in infection-induced SG formation among the three variants were due to differences in ISR activation, we next analyzed levels of PKR and eIF2α phosphorylation in infected A549-hACE2 cells. Clear differences were noted for the three variants. Levels of p-elF2a were highest in omicron-infected cells, and higher levels were observed in cells infected with the ancestral SARS-CoV-2 compared to Delta infected cells (Figure 1D and E). Omicron and the ancestral strain induced similar levels of p-PKR while the levels were clearly lower in Delta-infected cells (Fig. 3D and 3E).

### Effects of stress inducers on infection rate and progeny virus production for ancestral SARS-CoV-2, Delta, and Omicron

After observing differences in ISR activation and SG formation between the SARS-CoV-2 variants, we next compared their capacity to withstand stress stimuli. To this end, we analyzed the infection rate of the ancestral strain and the Delta and Omicron variants in A549-hACE2 cells treated with different stress inducers concurrently with the infection (Fig. 4A). As observed in ancestral strain-infected Vero E6 cells (Fig 2E), thapsigargin treatment almost completely blocked infection with all three variants in A549-hACE2 cells (Fig. 4A). In A549-hACE2 cells, significant reduction in infectivity was also observed with Poly(I:C) treatment for all variants and SA treatment for the ancestral strain and Delta, while the effect of starvation was small to negligible (Fig 4A). Analysis of viral titers following post treatments with stress inducers (added to the cells at 18 hpi) showed resistance to Poly(I:C) and susceptibility to thapsigargin by all three variants, but responses to SA and starvation differ between Omicron and the other two variants (Fig 4B).

**Figure 4:**
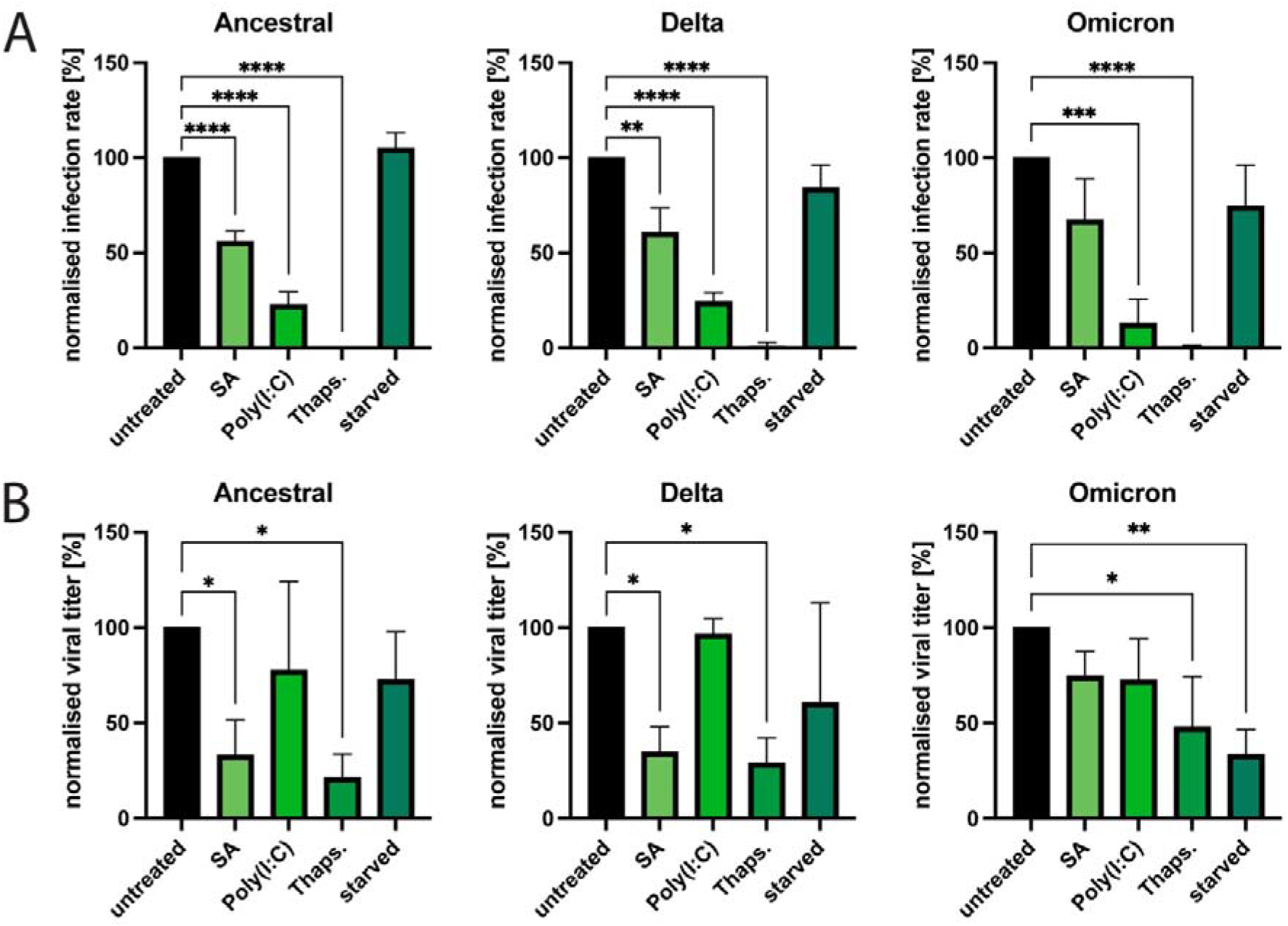
The effect of different stress stimuli on infectivity and replication of ancestral SARS-CoV-2, Delta, and Omicron. A) Effect of different stress inducers on infectivity. SARS-CoV-2 infected A549-hACE2 cells were treated with sodium arsenite, Poly(I:C), thapsigargin or starved as described in figure 2 and fixed at 24 hpi. They were stained for SARS-CoV-2 N protein. For the analysis using fluorescence microscopy, ≥5 images of a total of >100 cells were taken at random positions and the number of infected cells was determined and normalized against the infection rate of untreated cells. Data are represented as mean ± SEM. n = 3. (B) Effect of different stress inducers on viral replication. SARS-CoV-2 infected A549-hACE2 cells were treated with sodium arsenite, Poly(I:C), thapsigargin or starved as described in figure 2. 6h after treatment the viral titers were determined and normalized against the titers in untreated cells. Data are represented as mean ± SEM. n = 3. *, p < 0.05; **, p < 0.005; ***, p < 0.0005; ****, p < 0.0001

### Activation of the ISR contributes to translational arrest of infected cells and is beneficial for SARS-CoV-2 replication

During activation of the ISR, eIF2α-phosphorylation leads to a global translation arrest [11]. To assess whether the phosphorylation of eIF2α observed in SARS-CoV-2 infected cells (Fig. 1A and Fig. 3D) triggers a translational shutdown, infected A549-hACE2 cells were exposed to puromycin followed by analysis of total levels of newly produced proteins (Fig. 5A and 5B). At 24h after infection with ancestral SARS-CoV-2, Delta, or Omicron, global host cell protein production was clearly inhibited (Fig 5A and 5B). Treatment with sodium arsenite reduced host cell protein production even more (Fig 5A and 5B), suggesting that despite already high phosphorylation of eIF2α infected cells remain susceptible to increased translational arrest mediated by additional ISR activation. Further analysis by immunofluorescence microscopy showed that translational levels were reduced as early as 6 hpi in infected Vero E6 cells (suppl. Fig 3A and 3B).

**Figure 5:**
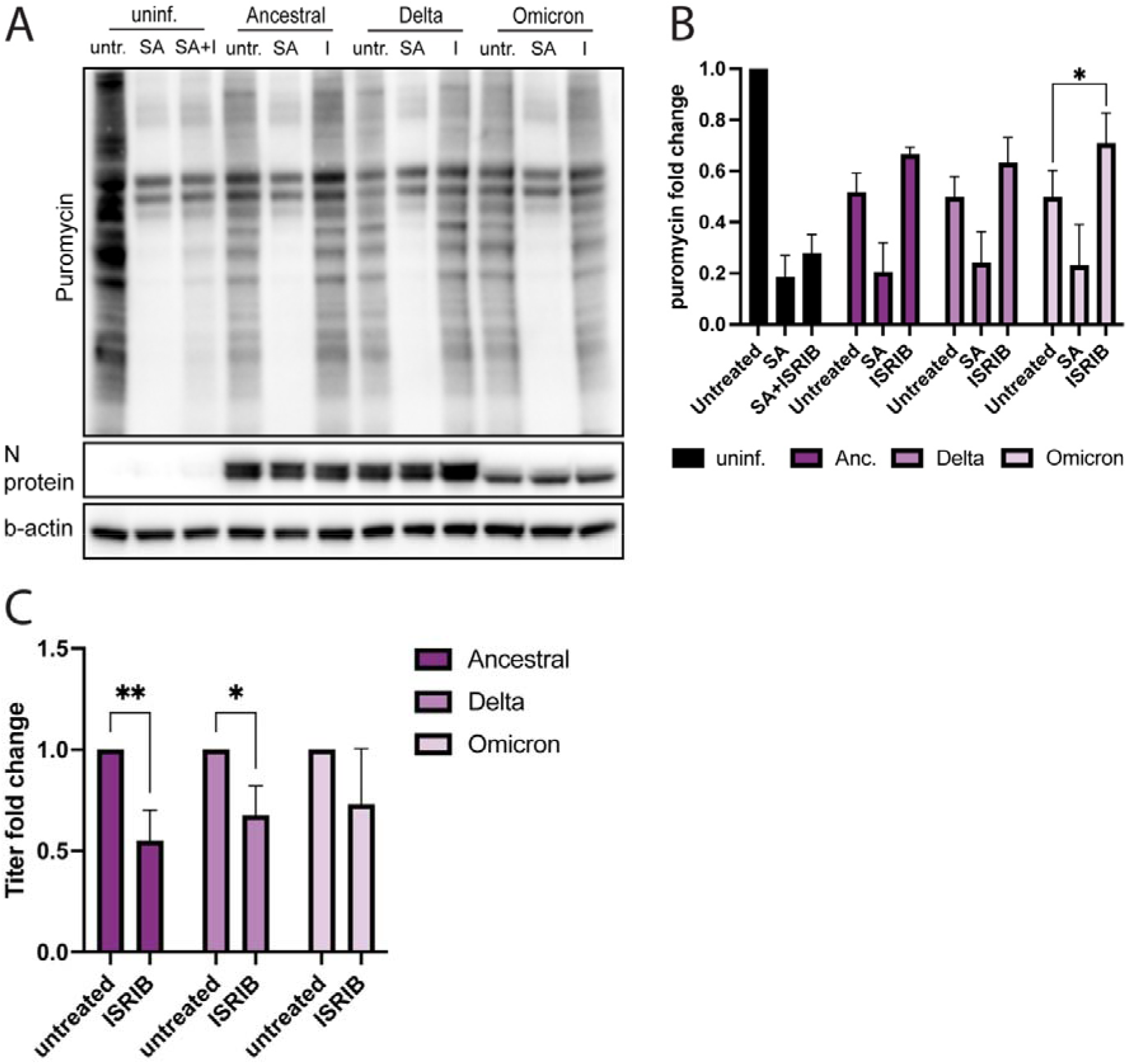
The effect of ISR activation during SARS-CoV-2 infection on global translational levels and viral replication in A549-hACE2 cells. (A) Puromycin incorporation in A549-hACE2 cells infected with SARS-CoV-2 variants. Uninfected and infected cells were treated with sodium arsenite and/or ISRIB. Before sample collection at 24 hpi, the cells were treated with puromycin to visualize translational levels. (B) Quantification of (A). Puromycin levels were normalized to cellular levels of β-actin. The data are presented as the fold change in relation to the puromycin levels of untreated, uninfected cells and are represented as mean ± SEM. n ≥ 3. (C) ISRIB treatment decreases viral replication. SARS-CoV-2 infected cells were treated with ISRIB, and the viral titers of the supernatant collected at 24 hpi was determined. The titers have been normalized against the titer of untreated cells and the data are represented as mean ± SEM. n = 3. *, p < 0.05; **, p < 0.005

To further investigate the extent to which ISR activation contributes to the translational arrest after SARS-CoV-2 infection (Figure 5A), infected cells were treated with the ISR inhibitor ISRIB, which blocks the interaction between p-eIF2α and eIF2B, thereby inhibiting p-eIF2α-mediated translational shutdown [30]. ISRIB treatment restored translation in infected cells to some extent (Figure 5A and 5B, Suppl. Fig. 4A and 4B), but the impact was significant only in omicron-infected cells (Figure 5A and 5B). ISRIB treatment also reduced SG formation at 6 hpi (Suppl Fig. 4C), indicating that the ISR plays a role in formation of SGs and triggering translational shutdown at early stages of infection. To analyze the impact of ISR activation on viral replication for the three variants, we compared viral titers of untreated and ISRIB treated SARS-CoV-2 infected A549-hACE2 cells (Fig. 5C). Overall, inhibiting the ISR had a negative effect on the replication of all three variants, with significantly lower titers observed in ancestral strain and Delta infected cells. Intriguingly, this indicates that the translational inhibition resulting from ISR activation during SARS-CoV-2 infection might be beneficial for the virus production.

### Expression of ATF4 and CHOP is suppressed in SARS-CoV-2 infected cells

While phosphorylation of eIF2α leads to a reduction in global translational levels, selected stress-responsive transcripts are still translated and even upregulated under these conditions [11]. This includes the transcription factor ATF4, a master regulator that controls the expression of a variety of proteins that in turn enable the cell to recover from stress, promote cell survival and eventually terminate the ISR via dephosphorylation of eIF2α [31]. However, if the stress is too intense or long-lasting, ATF4 can also induce apoptosis, for example through upregulation of the transcription factor CHOP [32-33]. Since SARS-CoV-2 infection induced eIF2α-phosphorylation (Figure 1A), we analyzed whether this also triggered the expression of ATF4 and its downstream effector CHOP. Interestingly, neither ATF4 nor CHOP could be detected in ancestral strain-infected A549-hACE2 cells and even SA-treatment failed to induce ATF4 production in infected cells (Figure 6A). Treatment with thapsigargin induced strong expression of ATF4 and CHOP in uninfected and infected cells, but the levels of both were clearly reduced in infected cells despite high p-eIF2α levels, suggesting a viral mechanism inhibiting the ISR downstream of eIF2α (Figure 6A).

**Figure 6:**
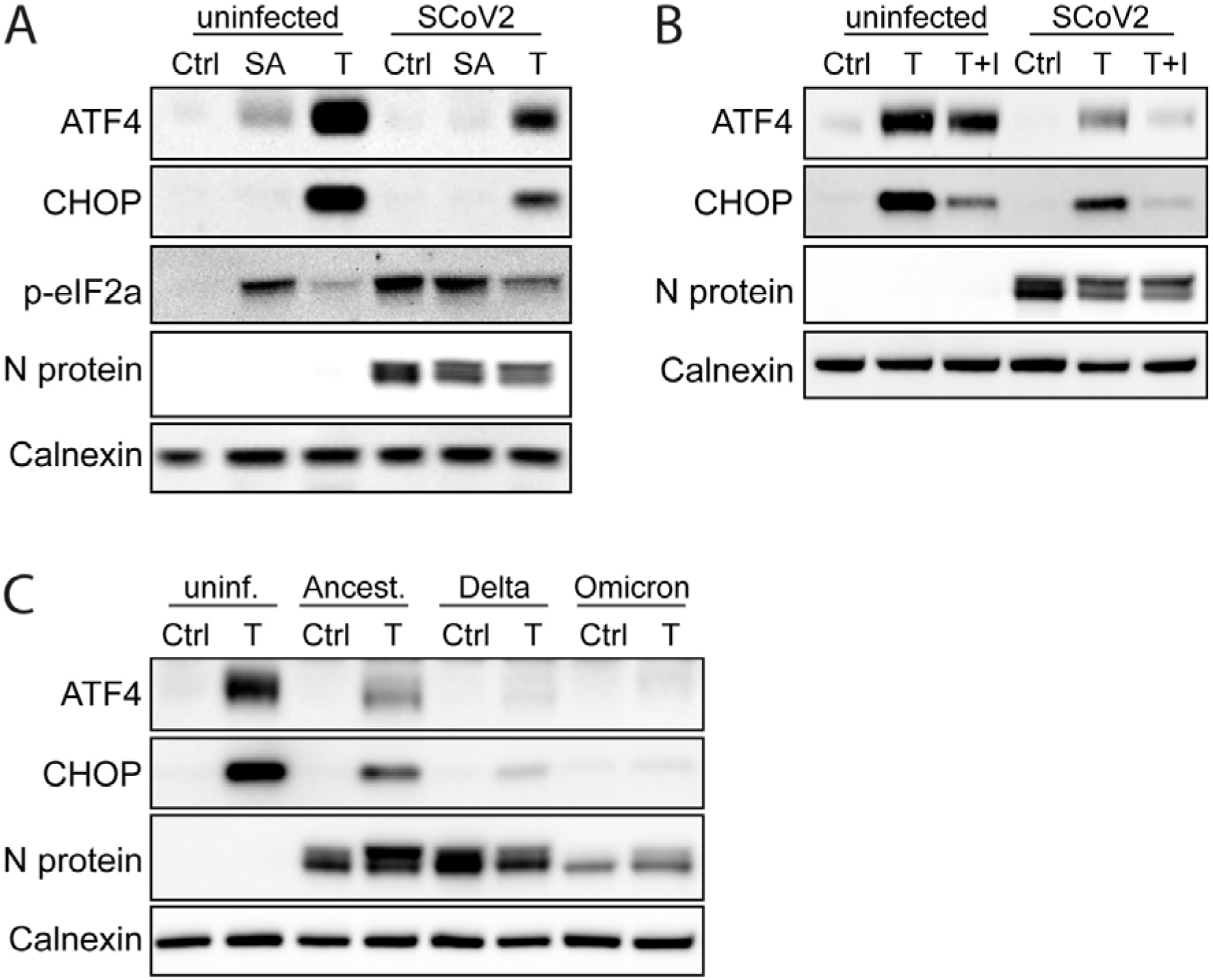
Suppression of ATF4 and CHOP production in SARS-CoV-2 infected A549-hACE2 cells. (A) Uninfected and ancestral strain-infected cells were treated with sodium arsenite (SA), thapsigargin (T) or remained untreated. At 24 hpi, cell lysates were collected and analyzed for expression of ATF4, CHOP and p-eIF2α via western blot. Calnexin was used as a loading control. (B) Uninfected and ancestral strain-infected cells were treated with T and ISRIB (I) or remained untreated. At 24 hpi, cell lysates were collected and analyzed for expression of ATF4 via western blot. Calnexin was used as a loading control. (C) Uninfected cells and cells infected with different SARS-CoV-2 variants were treated with T or remained untreated. At 24 hpi, cell lysates were collected and analyzed for expression of ATF4 via western blot. Calnexin was used as a loading control.

Interestingly, ISRIB treatment had a similar impact as SARS-CoV-2 infection on thapsigargin-induced production of ATF4 and CHOP, reducing the expression of both proteins in uninfected cells and further decreasing their levels in infected cells to an almost undetectable level (Figure 6B). Comparison of the SARS-CoV-2 variants showed even stronger inhibition of ATF4 and CHOP expression by Delta and Omicron variants, suggesting that ISR inhibition is a feature shared by all SARS-CoV-2 variants (Fig. 6C).

## Discussion

Viral infections disrupt cellular homeostasis, which in turn can trigger stress responses. Here we show that SARS-CoV-2, via activation of the eIF2α-kinase PKR, activates the ISR. When investigating downstream effectors of the ISR, we found that the production of ATF4 and CHOP is inhibited in infected cells. Further, despite strong eIF2α phosphorylation and a reduction in cellular translational levels, SG prevalence remains mostly low in infected cells. When comparing the ancestral SARS-CoV-2 and the two variants Delta and Omicron, we noted that Delta induces weaker levels of p-PKR and p-eIF2α, and Omicron induces stronger SG formation, suggesting variant specific differences.

A key player of the ISR is eIF2 which, when phosphorylated at Ser51 in its alpha subunit, induces an arrest in cap-dependent translation [34]. Four kinases, each activated by a specific type of stress, are known to phosphorylate eIF2α and three of them (PKR, PERK and GCN2) are known to be activated during viral infections [13]. Here, we observed strong phosphorylation of PKR, a kinase that detects double-stranded RNA and hence often is activated during viral infections [35]. While many viruses have developed strategies to block its function, we detected high levels of p-eIF2α in SARS-CoV-2 infected cells, indicating PKR-mediated activation of the ISR. This is in line with Li and colleagues, who reported phosphorylation of PKR and eIF2α in infected cells [27]. SARS-CoV-1 mediated PKR and eIF2α phosphorylation have been described by Krähling and colleagues [28]. For SARS-CoV-1, it was shown that in addition to PKR, PERK significantly contributes to eIF2α phosphorylation in infected cells [28]. Coronavirus glycoproteins can induce rearrangements of the ER membrane, forming double-membraned vesicles, inducing ER stress and subsequent PERK activation [36-37]. While we did not observe PERK phosphorylation at the time of sample collection, another study did report PERK activation during early SARS-CoV-2 infection [38]. Another parallel to SARS-CoV-1 [28] is the finding of SARS-CoV-2 mediated downregulation of GCN2. GCN2 was recently identified as an antiviral factor that sensitized host cells for SARS-CoV-2 induced cell death [39]. Hence, SARS-CoV-2 mediated downregulation of GCN2 might potentially counteract the antiviral properties of this kinase. Phosphorylation of eIF2α and the subsequent reduction in global translational levels normally trigger the formation of SGs. Nevertheless, despite high levels of p-eIF2α and a strong translational shutdown, most cells infected with SARS-CoV-2 remained negative for SGs, showing that SARS-CoV-2 counteracts this cellular response. The few SG-positive cells detected showed a common pattern of low N protein levels, in line with previous findings that the SARS-CoV-2 N protein localizes to SGs through liquid-liquid phase separation and can dissolve SGs by binding to G3BP [23, 40]. This may also explain why most SGs were found during early ancestral SARS-CoV-2 infection before sufficient amounts of N protein have been produced.

We observed that SARS-CoV-2 efficiently suppressed SG formation when stimulated with sodium arsenite or Poly(I:C), but upon challenge with ER stress or starvation infected cells showed an increased susceptibility towards SG formation. Why SGs form more frequently during ER stress and starvation is unclear but since both types of stress affect the cellular protein turnover, they might also influence N protein production or modification and therefore its capability to interact with G3BP. High levels of ATF4, the effector that regulates eIF2α dephosphorylation via production of GADD34, may partly explain why uninfected cells are free of SGs after treatment with thapsigargin.

As shown previously [41], we noted that thapsigargin-treatment efficiently inhibited SARS-CoV-2 infection. Furthermore, both thapsigargin and sodium arsenite had a strong negative effect on viral replication in A549-hACE2 cells. In contrast, Poly(I:C), while reducing the infection rate, barely affected progeny virus production. Poly(I:C) mimics viral infection and triggers PKR activation. Since PKR is already highly activated during SARS-CoV-2 infection, additional Poly(I:C) treatment might not be able to activate PKR any further and the virus likely has developed mechanisms to cope with this kinase’s antiviral effects. Thapsigargin and sodium arsenite on the other hand also trigger additional pathways besides the ISR that can negatively affect viral replication including the unfolded protein response and mTOR signaling [42-43], which may explain why the virus is more sensitive towards these treatments.

An analysis of the translational activity during late-stage SARS-CoV-2 infection revealed a strong suppression of translation in infected cells. While the high levels of p-eIF2α present during the infection might suggest ISR activation as the mechanism behind the translational arrest, the fact that ISRIB treatment largely failed to restore translation indicated it is not. Instead, the bulk of the translational arrest might be mediated by nsp1, an important SARS-CoV-2 virulence factor known to suppress host gene expression by binding to the entry channel of ribosomes and blocking translation of host mRNAs while favoring viral mRNAs [44–46]. Worth noting is, however, that ISRIB treatment reduced SG formation at early time points of SARS-CoV-2 infection, indicating that ISR activation plays a more significant role in triggering translational arrest early during the infection. It is also noticeable that while p-eIF2α levels of uninfected, sodium arsenite-treated A549-hACE2-cells and SARS-CoV-2 infected cells were similar, sodium arsenite had a stronger effect on translational levels than infection *per se*.

During ISR activation, the binding of p-eIF2α to eIF2B leads to a reduction in availability of the eIF2-GTP-tRNAiMet-ternary complex which is an essential factor for translation initiation [47–49]. It has been recently shown that the Beluga whale coronavirus SW1 employs a class IV ISR inhibitor which blocks the interaction between p-eIF2α and eIF2B to avert translational arrest in the presence of high p-eIF2α concentrations [50-51]. Possibly, SARS-CoV-2 could similarly utilize a class IV ISR inhibitor to assure the availability of sufficient ternary complex levels for its own protein production. In line with this, we found that levels of a key regulator of the ISR, the transcription factor ATF4, as well as of its effector protein CHOP, were both suppressed in SARS-CoV-2 infected cells. Both proteins are normally upregulated and preferentially translated during p-eIF2α mediated translational arrest [11]. Under prolonged or intensive stress conditions ATF4 and CHOP promote apoptosis [33]. Apoptosis is a common mechanism for elimination of infected cells. Consequently, many viruses have developed anti-apoptosis mechanisms, for instance upregulating the anti-apoptotic factor BCL-2 and blocking of DR5 expression of [52-53]. Employing a class IV ISR inhibitor could therefore be a strategy to simultaneously avoid a lack of ternary complex as well as avert apoptosis during persistent ISR signaling. Since ISR activation is one of the host cells’ ways to fight off viral infections, it is perhaps surprising that inhibition of the pathway did not enhance SARS-CoV-2 replication but instead lead to a decrease in viral titers. This suggests that SARS-CoV-2 infection already copes so well with ISR activation that inhibiting the pathway actually hampers viral replication conceivably by benefiting host cell translation more than viral translation.

When comparing ancestral SARS-CoV-2 with the Delta and Omicron variants, we observed several variant-specific differences in regard to ISR activation and SG formation. All variants triggered ISR activation, but Delta induced significantly less phosphorylation of PKR and eIF2α than the ancestral strain and Omicron. It has been suggested that the SARS-CoV-2 N protein can inhibit PKR activation by sequestering dsRNA [24, 54] and adaptations in the Delta N protein might make this process more efficient. Omicron on the other hand induces similar levels of p-PKR but increased levels of p-eIF2α, indicating stronger ISR activation. No other eIF2α-kinases than PKR were activated by ancestral SARS-CoV-2 but PERK or GCN2 might play a role in ISR activation during Omicron infection and trigger additional eIF2α phosphorylation. The high concentrations of p-eIF2α could partly explain why Omicron responds differently to ISR inhibition. ISRIB treatment restored translational levels more efficiently in Omicron infected cells than in cells infected with ancestral SARS-CoV-2 or Delta, suggesting that the ISR contributes more to inducing translational arrest than with the other variants. In contrast to ancestral SARS-CoV-2 and the Delta variant, Omicron strongly induced SG formation, especially at late time points of the infection. The elevated levels of p-eIF2α in Omicron-infected cells could contribute to this effect. Intriguingly, unlike in ancestral SARS-CoV-2 or Delta-infected cells, almost no difference in N protein levels could be observed between Omicron-infected SG-negative and SG-positive cells. This suggests that the Omicron N protein might be less efficient in SG suppression than the N proteins of the other variants. The Omicron N protein contains several mutations within the intrinsically disordered region of its N-terminus, which contributes to the protein’s ability to induce phase separation, the process through which the N protein likely localizes to SGs [55]. Further supporting the possible involvement of the N protein, Barh and colleagues recently reported that the binding of the Omicron N protein to G3BP is weaker than that of ancestral SARS-CoV-2 or Delta N proteins [56].

To summarize, we report that while SARS-CoV-2 strongly activates the ISR through PKR leading to phosphorylation of eIF2α, the prevalence of SGs in infected cells remains low and no expression of stress-responsive transcripts is observed. Treatments with ISRIB and different stress kinase inducers shows SARS-CoV-2 resistance especially towards PKR activation and indicates a balance where ISR activation resulting from SARS-CoV-2 infection appears beneficial for the virus possibly through the hindrance it causes on host translation, but overt stimulation of the stress kinases will result in inhibition of viral replication. Finally, we show variant-specific differences in SARS-CoV-2 activation of the ISR, particularly in that the Omicron variant causes significantly more formation of stress granules compared to the ancestral strain and Delta.

## Acknowledgements

We thank Dr Pamela Österlund from the National Institute for Health and Welfare, Finland, for providing us with the SARS-CoV N antibody and Oscar Fernandez-Capetillo’s lab at the Science for Life Laboratory in Stockholm for providing us with the A549-hACE2 cells.

This study was supported by grants from CIMED and SciLifeLab/KWA.

## Materials and Methods

### Cell culture

Green monkey kidney Vero E6 cells were grown in minimum essential medium (MEM) supplemented with 7.5% FBS, HEPES, L-glutamin, 100 U/ml penicillin, and 100 μg/ml streptomycin. hACE2-overexpressing human lung epithelial A549-hACE2 cells (a kind gift from Oscar Fernandez-Capetillo’s lab) were grown in MEM supplemented with 7.5% FBS, HEPES, L-glutamin, 100 U/ml penicillin, and 100 μg/ml streptomycin. Both cell types were grown at 37°C and 5% CO_2_.

### Viruses and infection

The ancestral SARS-CoV-2 (isolate SARS-CoV-2/human/SWE/01/2020; Genbank accession: MT093571), Delta and Omicron BA.1 strains were propagated on Vero E6 cells and titrated via end point dilution assay as described earlier (see below). Cells were infected in complete MEM at a multiplicity of infection (MOI) of 0.1. After one hour of incubation the virus solution was removed, and fresh growth medium was added. Samples were collected at the indicated time points.

### Immunofluorescence microscopy

Cells were fixed for 20 min with 4% (vol/vol) paraformaldehyde in PBS. Before the staining, the cells were permeabilized by treatment with 0.5% (vol/vol) triton X in PBS for 5 min and blocked with 0.5% (wt/vol) bovine serum albumin (BSA) in PBS for 30 min. Primary antibodies against the SARS nucleocapsid protein [57] (rabbit, 1:200), G3BP (abcam, ab181149) (mouse, 1:500) and puromycin (Merck, MABE343) (mouse, 1:500) were diluted in the blocking solution and incubated on the cells for 1h. After washing, the samples were incubated for 1h with the respective secondary antibodies, goat anti-mouse alexa-488 or goat anti-rabbit alexa-647, and DAPI diluted 1:1000 in blocking solution. Imaging was done on a Nikon A1R+ confocal microscope with NIS-Elements C software. The images are representative images from at least three independent experiments. For the manual quantifications, 100 to 200 cells on 5 to 10 random microscope frames were counted for each biological replicate. For the quantification of SGs in infected cells, only cells that were positive for the N protein were included in the counting. Measuring of N protein intensity by calculating the CTCF was performed with Fiji/ImageJ version 2.0.0 software. The acquisition settings were kept constant between all conditions to ensure comparability.

### Western blotting

Cells were washed with PBS and lysed with 100 μL RIPA buffer (Thermo Scientific) supplemented with protease inhibitors (Complete; Roche) and phosphatase inhibitors (PhosStop, Sigma). NuPage sample buffer with 1.5% beta-mercaptoethanol was added to the samples before incubating them for 10 min at 96°C. The proteins were separated via SDS-PAGE (10% or 4-12% polyacrylamide gel) and transferred to a polyvinylidene difluoride (PVDF) membrane with an iBlot 2 dry blotting system. The membranes were then blocked with 5% (wt/vol) milk powder in Tris-buffered saline (TBS) with 0.1% Tween (TBS-T) for 1h. The following primary antibodies were diluted in 5% (wt/vol) BSA–TBS-T and incubated at 4°C overnight: rabbit anti SARS nucleocapsid protein [57] (1:2500), rabbit anti-phospho-eIF2α (cell signaling, #9721) (1:1000), rabbit anti-phospho-PKR (abcam; ab32036) (1:1000), rabbit anti-PKR (cell signaling, #12297) (1:1000), rabbit anti-phospho-PERK (Thermo Fisher, PA5-40294) (1:500), rabbit anti-PERK (cell signaling, #5683) (1:1000), rabbit anti-phospho-GCN2 (abcam, ab75836) (1:1000), rabbit anti-GCN2 (cell signaling; #3302) (1:1000), mouse anti-puromycin (Merck, MABE343) (1:10.000), rabbit anti-ATF4 (cell signaling, #11815) (1:1000) and mouse anti-CHOP (cell signaling, #2895). The following antibodies were diluted in 5% (wt/vol) milk powder–TBS-T and incubated at room temperature for 1h: rabbit anti-calnexin (cell signaling; #2679) (1:1000) and mouse anti-beta-actin (cell signaling; #3700) (1:10,000). The secondary, horseradish peroxidase (HRP)-conjugated goat antibodies were diluted 1:10,000 in 5% (wt/vol) milk powder in TBS-T and incubated for 1 h at room temperature. Images were obtained with GelCapture (DNR Bio-Imaging Systems). The images shown are representative of results from at least three independent experiments. Densitometric quantification of bands was done with Fiji/ImageJ version 2.0.0 software.

### Treatments

To induce stress and SG formation, the cells were treated with 1mM sodium arsenite (sigma) for 1 h, 5μM thapsigargin (Merck; T9033) for 6 h, HBSS (+2% HEPES + 2% FBS) for 6 h to starve the cells. For treatment with Poly(I:C), cells were transfected with 2.5μg Poly(I:C) using Lipofectamine LTX with PLUS reagent (Thermo Fisher, 15338100). When analyzing the effect of different stress inducers on viral replication, the cells were treated with 0.1mM sodium arsenite, 5μM thapsigargin or HBSS (+2% HEPES + 2% FBS) for 6 h. When analyzing the effect of different stress inducers on infectivity, the cells were treated with 10 μM sodium arsenite, 0.5μM thapsigargin or medium containing 50% HBSS (+2% HEPES + 2% FBS) for 24 h or transfected with 0.25μg Poly(I:C). For the puromycin labeling, the cells were treated with 20μM puromycin for 10 min and washed twice with PBS before sample collection. Infected cells were treated with 200nM ISRIB (Merck; SML0843) throughout the infection. For uninfected cells, 200nM ISRIB was added together with the sodium arsenite.

### End point dilution assay

Vero E6 cells in 96-well plates were treated with supernatants from infected cells in a dilution series of 10^−1^ to 10^−8^ with eight wells per dilution. After 5 days the cells were analyzed for cytopathic effect (CPE) with a light microscope to detect infection. Wells that were positive for CPE were marked and the number of 50% tissue culture infectious doses (TCID_50_) per milliliter was calculated using the Spearman-Karber method.

### Statistical analysis

Error bars show standard deviations. t tests (Fig. 1 and suppl. Fig 4C), one-way analysis of variance (ANOVA) (Fig. 3B and 3E, Figure 4A and 4B) and two-way ANOVA (Fig. 2B-G, Fig. 3A and 3C, Fig. 5B and 5C, suppl. Fig 2B and suppl. Fig 4B) were performed in GraphPad Prism Version 9.4.0 (453) to calculate P values (*, p < 0.05; **, p < 0.005; ***, p < 0.0005; ****, p < 0.0001).

---

**Supplementary Figure 1:**
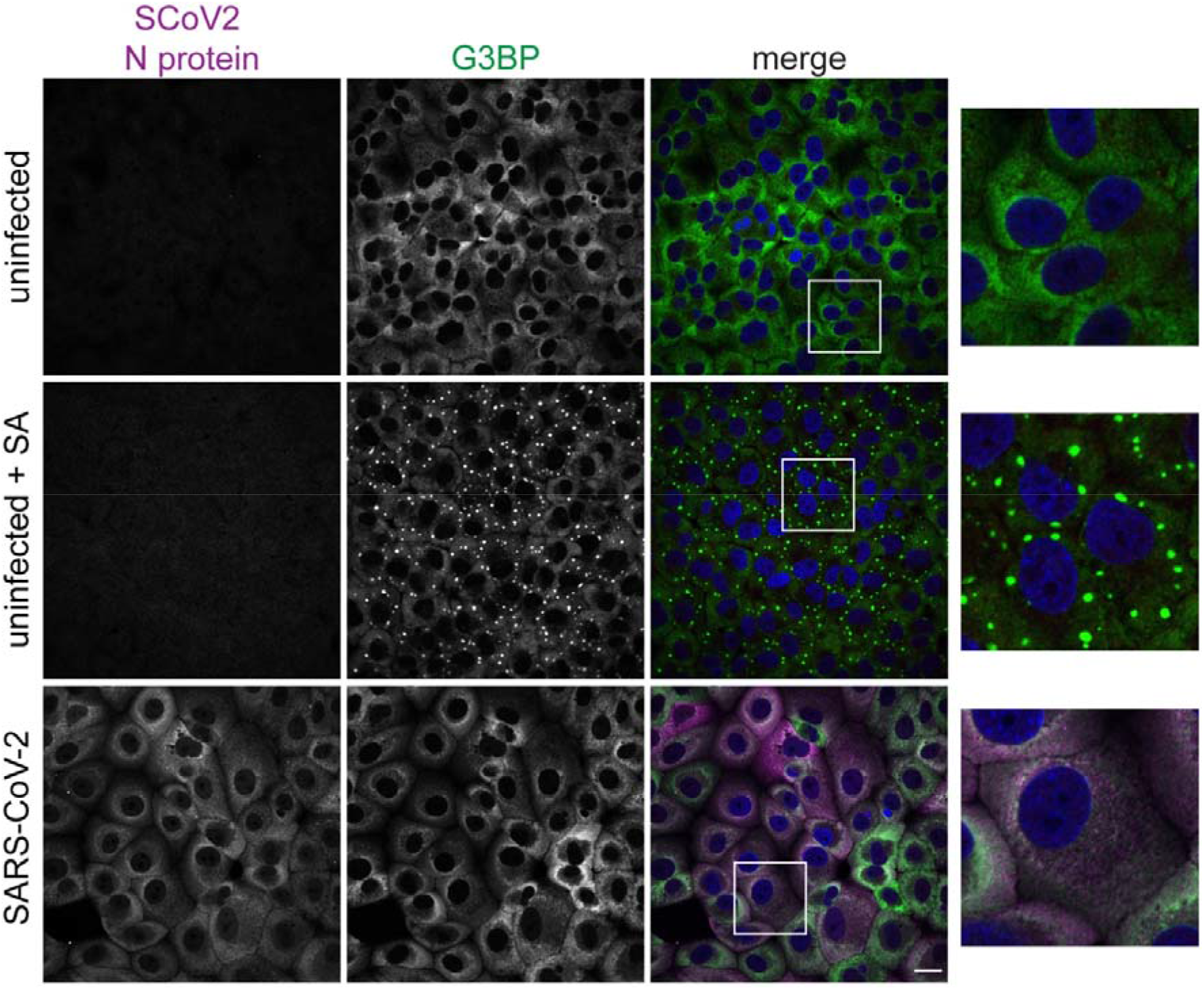
G3BP staining in ancestral SARS-CoV-2 infected Vero E6 cells. Cells were infected with ancestral SARS-CoV-2 and fixed after 24h. Uninfected cells that were either untreated or treated with 1mM sodium arsenite (SA) were used as a control. The cells were stained with antibodies against SARS-CoV-2 N protein (magenta) and G3BP (green) and with DAPI (blue). Imaging was performed with confocal microscopy (20X). Scale bars 20 μm.

**Supplementary figure 2:**
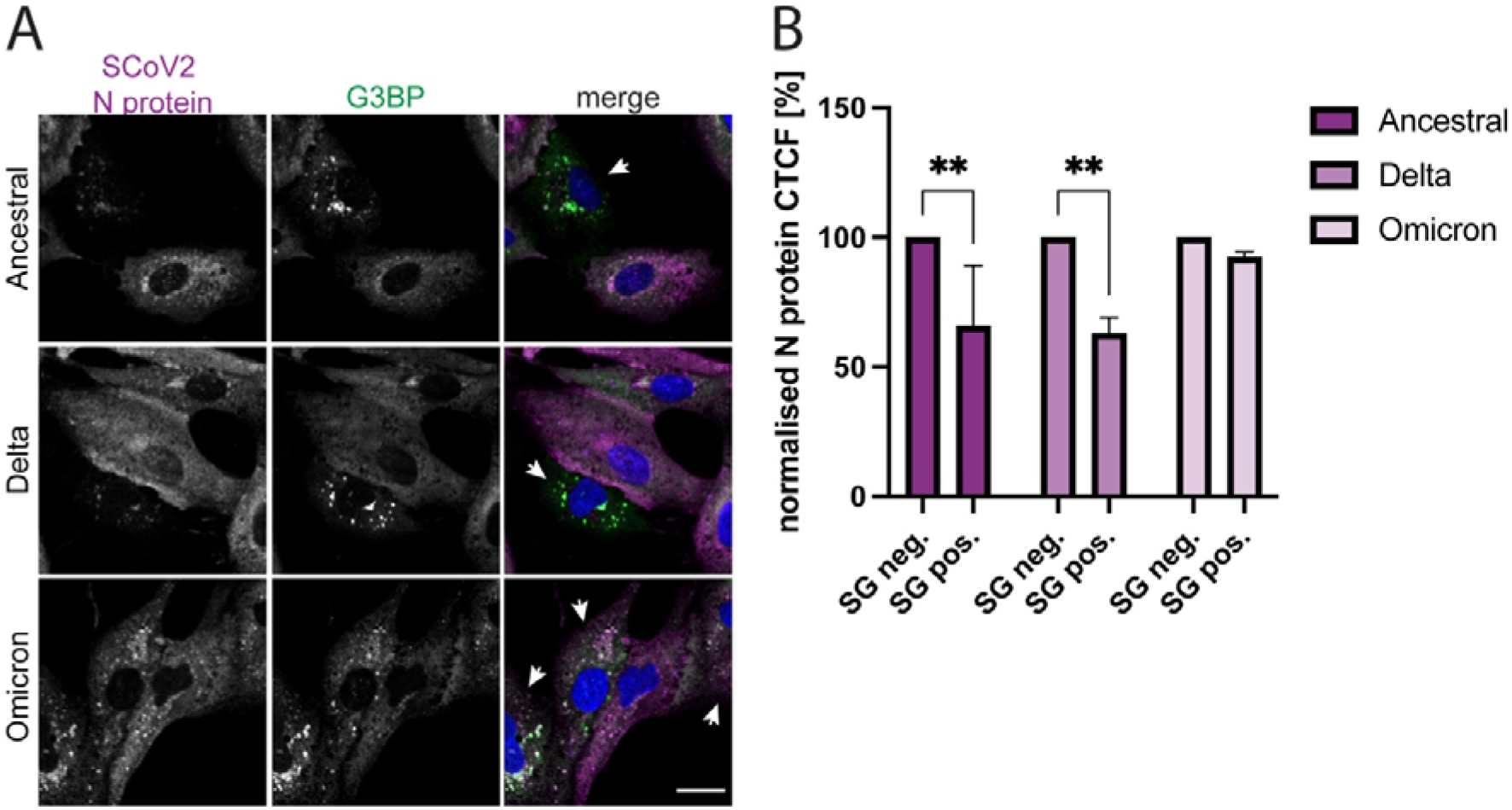
Nucleocapsid protein levels in SG-negative and -positive A549-hACE2 cells infected with different SARS-CoV-2 variants. (A) N protein levels in individual A549-hACE2 cells. Ancestral SARS-CoV-2, Delta, or Omicron infected A549-hACE2 cells were fixed 24 hpi and stained with antibodies against SARS-CoV-2 N protein (magenta) and G3BP (green) and with DAPI (blue). Arrows show SG-positive cells. Imaging was performed with confocal microscopy (20X). Scale bars 20 μm. (B) Quantification of N protein levels in SG-negative and -positive A549-hACE2 cells. Ancestral SARS-CoV-2, Delta, or Omicron infected A549-hACE2 cells were fixed 24 hpi and the N protein corrected total cell fluorescence (CTCF) of SG-negative and SG-positive cells was determined. For each variant, the CTCF of SG-positive cells was normalized against the CTCF of SG-negative cells. Data are represented as mean ± SEM. n = 3 **, p < 0.005

**Supplementary Figure 3:**
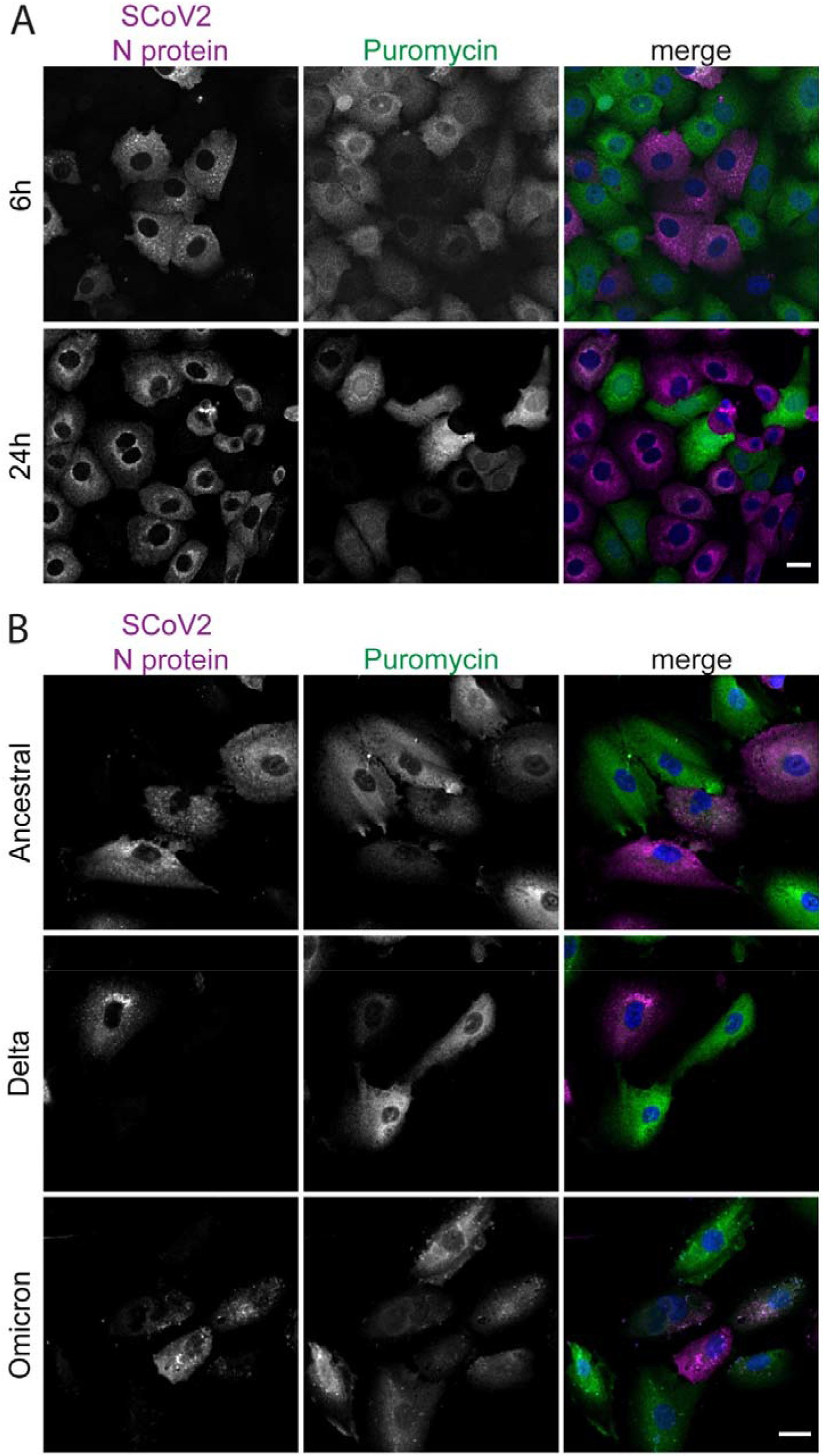
Translational levels in ancestral SARS-CoV-2 infected cells. (A) Vero E6 cells were infected with ancestral SARS-CoV-2. Before fixation at 6 or 24 hpi, the cells were treated with puromycin. The cells were stained with antibodies against the viral N protein (magenta) and puromycin (green), and DAPI (blue). Scale bars 20 μm. (B) A549-hACE2 cells were infected with different SARS-CoV-2 variants. Before fixation at 24 hpi, the cells were treated with puromycin. The cells were stained with antibodies against the viral N protein (magenta) and puromycin (green), and DAPI (blue). Scale bars 20 μm.

**Supplementary Figure 4:**
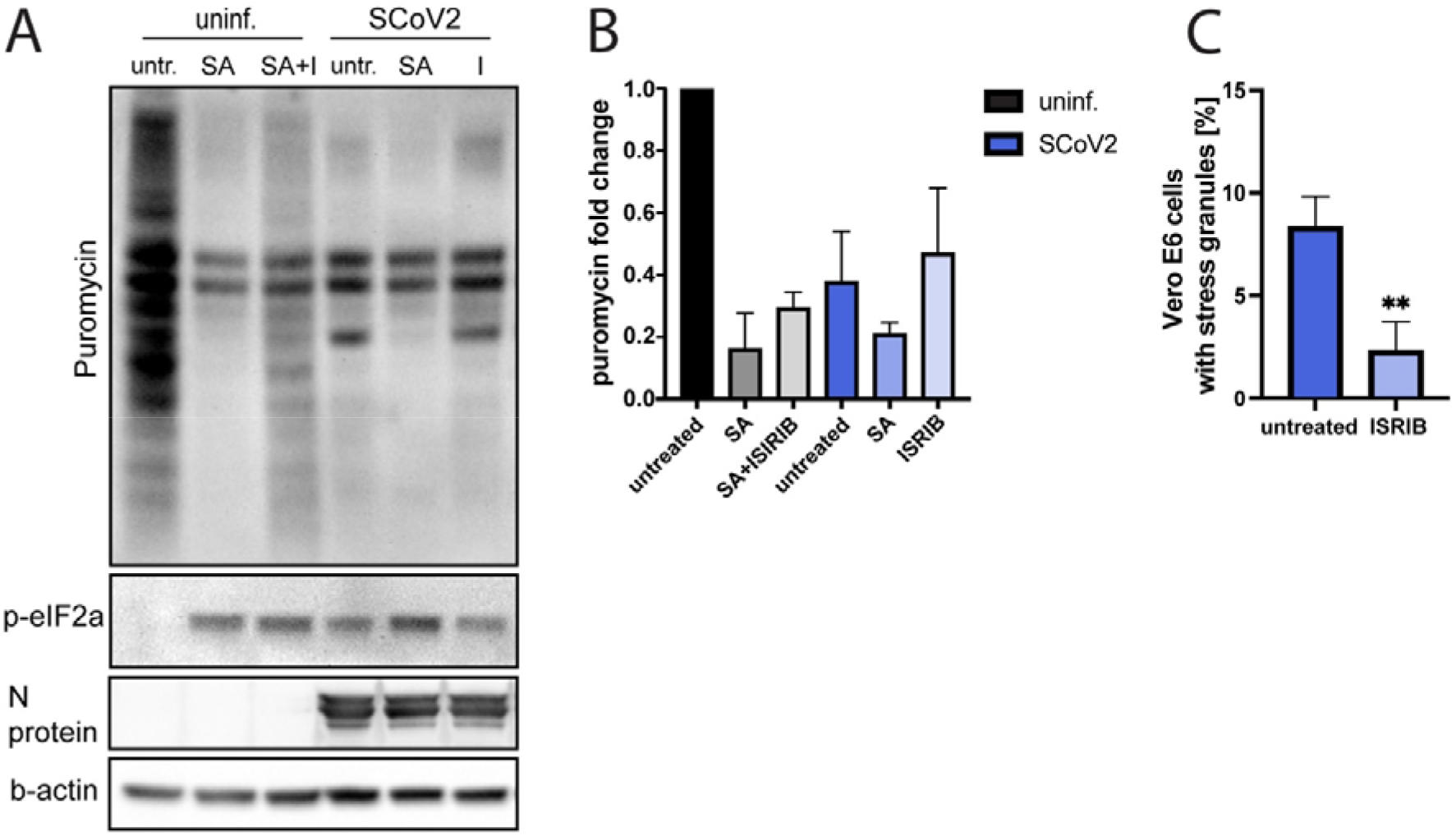
The effect of ISR activation during SARS-CoV-2 infection on global translational levels and SG formation. (A) Puromycin incorporation in ancestral SARS-CoV-2 infected Vero E6 cells. Uninfected and infected cells were treated with sodium arsenite (SA) and/or ISRIB (I). Before sample collection at 24 hpi, the cells were treated with puromycin to visualize translational levels. (B) Quantification of (A). Puromycin levels were normalized to cellular levels of β-actin. The data are presented as the fold change in relation to the puromycin levels of untreated, uninfected cells. Data are represented as mean ± SEM. n ≥ 3. (C) ISRIB treatment reduces SG formation. Ancestral strain-infected Vero E6 cells were treated with ISRIB and fixed at 6 hpi. They were stained for SARS-CoV-2 N protein and G3BP. For the analysis using fluorescence microscopy, ≥5 images of a total of >100 cells were taken at random positions and the number of infected cells showing SGs was determined. Data are represented as mean ± SEM. n = 3. *, p < 0.05; **, p < 0.005

